# Density Dependent Resource Budget Model for Alternate Bearing

**DOI:** 10.1101/2020.06.15.148924

**Authors:** Shadisadat Esmaeili, Alan Hastings, Karen Abbott, Jonathan Machta, Vahini Reddy Nareddy

## Abstract

Alternate bearing, seen in many types of plants, is the variable yield with a strongly biennial pattern. In this paper, we introduce a new model for alternate bearing behavior. Similar to the well-known Resource Budget Model, our model is based on the balance between photosynthesis (carbon accumulation) and reproduction processes. We consider two novel features with our model, 1) the existence of a finite capacity in the tree’s resource reservoir and 2) the possibility of having low (but non-zero) yield when the tree’s energy level is low. We achieve the former using a density dependent resource accumulation function, and the latter by removing the concept of the well-defined threshold used in the Resource Budget Model. At the level of an individual tree, our model has a stable two-cycle solution, which is suitable to model plants in which the alternate bearing behavior is pronounced. We incorporate environmental stochasticity by adding two uncorrelated noise terms to the parameters of the model associated with the carbon accumulation and reproduction processes. Furthermore, we examine the model’s behavior on a system of two coupled trees with direct coupling. Unlike the coupled Resource Budget Model, for which the only stable solution is the out-of-phase solution, our model with diffusive coupling has stable in-phase period-2 solutions. This suggests that our model might serve to explain spatial synchrony on a larger scale.

## 1. Introduction

“Alternate bearing” is the variability of fruit or nut production in many types of plants for which a year of high yield (ON-year) is followed by one or more years of low or no production (OFF-years). Generally, the crop varies biennially. However, in some cases, it can show longer period cycles where multiple years of high or low yield happen consecutively (Monselise and Goldschmidt, 1982; Shalom et al., 2012). When this phenomenon is observed in collective synchrony among trees in orchards and natural forests, it is known as masting.

Alternation is very common and is observed in a variety of plants like citrus trees (Shalom et al., 2012), olive trees (Lavee, 2007), and pistachio trees (Lyles et al., 2015; Noble et al., 2018). These plants are different in many ways. The differences include time of flowering, dormancy, and duration of fruiting compared to vegetative growth (Monselise and Goldschmidt, 1982). The ubiquity of the phenomenon suggests that there is a common mechanism that explains the crop variability in a variety of plants.

Both exogenous conditions, like environmental triggers, and endogenous factors, like bud abscission, flowering inhibition by current fruits (Shalom et al., 2012), pollination, and fruit overload, have been considered as contributing factors to the alternate bearing phenomenon. The depletion of the carbohydrate (resource) level of the plant due to over-fruiting is considered the most common cause of the phenomenon (Monselise and Goldschmidt, 1982; Lavee, 2007). Isagi pioneered a simple model to explain the mechanism of variable acorn yield observed at the level of an individual tree (Isagi et al., 1997). His model, called the Resource Budget Model (RBM), is based on the dynamics of the tree’s energy resource (carbohydrate) which is accumulated as the result of photosynthesis and consumed during the flowering and nut production processes.

The Resource Budget Model, as originally proposed by Isagi et al. (1997) and expanded in Iwasa and Satake (2004), assumes the existence of a well-defined threshold for the tree’s resource levels below which the plant will not reproduce. This means that during an OFF-year, the tree has no yield. This assumption is appropriate for the plants like olive and citrus, for which there is zero or near-zero yield during an OFF-year, but represents other species with low but positive OFF-year yields less well. Once the resource level of the tree exceeds the threshold, flowering and nut production happens. Both flowering and nut production processes are costly and result in the depletion of the tree’s resource reservoir. The cost of flowering and nut production is assumed to be proportional to the amount of resources above the threshold with the depletion coefficient (the parameter of the model). The Resource Budget Model belongs to the category of tent maps for which there is no stable period-2 solution at the level of individual tree (except at the bifurcation point). Systems of two trees do have an in-phase period-2 solution, but it is only stable if the trees are coupled via indirect (mean-field) coupling, as with pollination, and not for trees coupled directly (diffusively) through local interactions like root grafting (Prasad et al., 2017). Together, these features make the Resource Budget Model a simple and successful model to explain the masting phenomenon in many plants for which there is zero or near-zero yield during an OFF-year, the plant goes through several OFF-years before having a year with high yield, and at a collective level, the plants can interact via pollination (the plants are monoecious). But the model needs to be modified if it is to be applied to the plants like pistachio that have a low yield, but not zero, during OFF-years, whose yield show a two-cycle behavior, and is dioecious, therefore, the interaction between the trees happens via diffusive coupling (root grafting). Lyles et al. modified the Resource Budget Model by removing the concept of threshold from the model and adding temporal stochasticity to the depletion coefficient and the pollen availability to achieve the variable and synchronized nut production of the trees (Lyles et al., 2015). However, this model, like the previous Resource Budget Models, does not show synchrony in trees with direct (diffusive) coupling.

In another attempt to predict the yield of citrus trees, Ye and Sakai suggested a more generalized version of the Resource Budget Model (Ye and Sakai, 2016). Motivated by the result of their analysis of field data collected from a citrus orchard in Japan (Ye et al., 2008), they added a vegetative growth factor to the Resource Budget Model to account for the role of new leaf growth in inhibition of fruit production. According to their model, the cost of new leaf growth is proportional to the empty portion of the resource tank. Also, the return map obtained from their experimental study showed, what they called, a “hump-shaped” curve similar to what is obtained from the logistic map. To reproduce the logistic-like return map, they replaced the linear relationship between the resource reserve level and flowering and fruiting cost in the original Resource Budget Model (see Sec. 2), with a nonlinear Ricker-type relationship (Ye and Sakai, 2016). By adding nonlinearity they modified the model to better reflect the yield dynamics of citrus trees. However, similar to the original Resource Budget Model, this model assumes the existence of a well-defined threshold below which no flowering or production happens, which does not reflect the low (but non-vanishing) yield of species, like pistachio, during OFF years.

Inspired by an existing data set collected from a pistachio orchard at the level of individual trees during a 6-year period (Lyles et al., 2015; Noble et al., 2018), we propose a different approach for modeling alternate bearing that accommodates low but non-vanishing yield during OFF-years. This is achieved by replacing the concept of a sharp cut-off for reproduction (represented by a threshold function) with a continuous function that accommodates the non-zero yield when the tree’s current energy level is low. Also, we take into account the fact that there is a maximum capacity for the plant to store photosynthate and therefore, its energy storage cannot grow indefinitely.

In Sec. 2 we briefly review the rules and the characteristics of the Resource Budget Model. In Sec. 3, we describe our new model of alternate bearing behavior for the trees with low yield during OFF-years. We analyze the model by performing a bifurcation analysis. Furthermore, we discuss and apply some necessary constraints on the model to make it biologically meaningful and applicable. In Sec. 4, we add stochasticity to the model to account for environmental variation. As the preliminary step to understand the collective behavior of the trees in an orchard or a natural forest, in Sec. 5, we study the dynamics of a two-tree system.

## 2. Background

First we describe the Resource Budget Model: every year, the resource level of an individual tree increases by a constant amount called *P*_*s*_. If the resource level exceeds a threshold, *L*_*T*_, the plant will flower and bear fruits/nuts which depletes the energy reservoir of the tree. The cost of flowering is assumed to be proportional to the excess amount of resources above the threshold with a positive constant *a*. The cost of fruit/nut production is also considered to be proportional to the cost of flowering. The Resource Budget Model is formulated as,

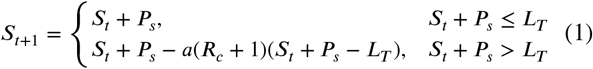

where *R*_*c*_ is the ratio of the cost of fruit/nut production to cost of flowering. The model can be written in terms of the dimensionless variable as, 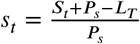 as,

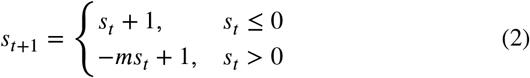

where *m* = *a*(*R*_*c*_ + 1) − 1 is called the depletion coefficient.

As it is shown in the model’s orbit diagram (Figure 1) and discussed in Appendix A, the model has a stable fixed point for *m* < 1. At exactly *m* = 1 (which is the bifurcation point) the system shows two-cycle behavior. For *m* > 1 the system demonstrates a chaotic period-four oscillation for a very small range of the parameter, followed by a single band chaos (Prasad and Sakai, 2015).

**Figure 1:**
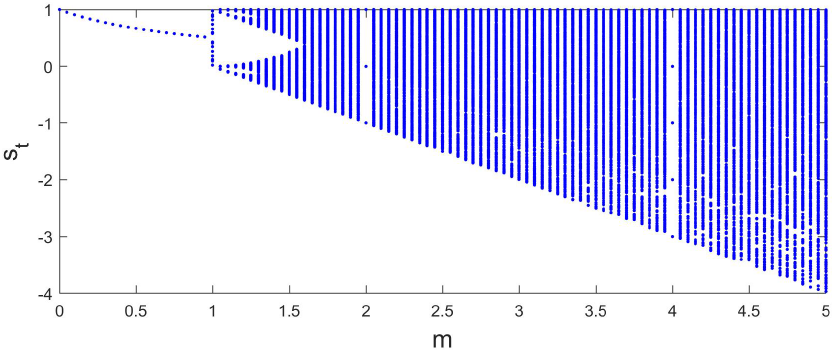
The orbit diagram of the Resource Budget Model shows that the dynamics of the system goes from a stable fixed point for the depletion coefficient, *m* < 1, to period-four oscillation for a very small range of *m* and then quickly leading to chaos.

## 3. The Model

Similar to the Resource Budget Model, our new model of alternate bearing is based on the dynamics of the ene (i.e. carbohydrate or carbon) level of an individual tree. According to our model, the carbon level of an individual tree in year *t* + 1 is determined based on the balance between two processes that happen in year *t*: 1) carbon accumulation, 2) reproduction. The accumulation of carbon is the result of photosynthesis (Marino et al., 2018). The process of reproduction is the flowering and production of nuts. Nut production comes with a higher cost than flowering and is considered the main sink of the tree’s carbon reservoir (Marino et al., 2018). The carbon level of a tree in year *t* + 1 (*S*_*t*+1_) can be written as,

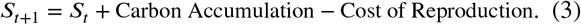

In modeling the Carbon Accumulation and the Cost of Reproduction, we take into account the following considerations:

1. The Carbon Accumulation process cannot result in the indefinite growth of the tree’s carbon level. In other words, each tree has a maximum capacity to store carbon, denoted by *S*_*max*_. Therefore, the amount of carbon that is added to the tree’s reservoir each year is a density dependent function of its existing carbon level. We model the Carbon Accumulation process by a function of *S*_*t*_, which grows when *S*_*t*_ is small but approaches zero as *S*_*t*_ → *S*_*max*_.
2. As we mentioned in the introduction, the function used to model the Cost of Reproduction should allow for low yield (as opposed to zero yield) during OFF-years.

There are many mathematical functions that satisfy the above conditions and can be considered to model these two processes. For the purpose of this paper, we have chosen the bounded growth function for Carbon Accumulation and a shifted sigmoid function for the Cost of Reproduction. However, these functions are not unique. In Appendix C, we present an alternate version of the model using a different Carbon Accumulation function and show that the dynamics of the model stay qualitatively similar.

### Bounded Growth Function

The amount of carbon added to the tree’s reservoir as a result of photosynthesis at the end of year *t* is modeled by:

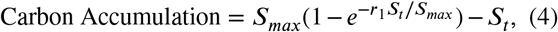

where *S*_*t*_ is the current carbon level, *S*_*max*_ is the tree’s maximum capacity to store carbon, and *r*_1_ is the efficiency of the Carbon Accumulation process. Figure 2a shows the behavior of the nondimensionalized version of equation 4 as a function of 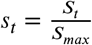 and for different values of *r*_1_

**Figure 2:**
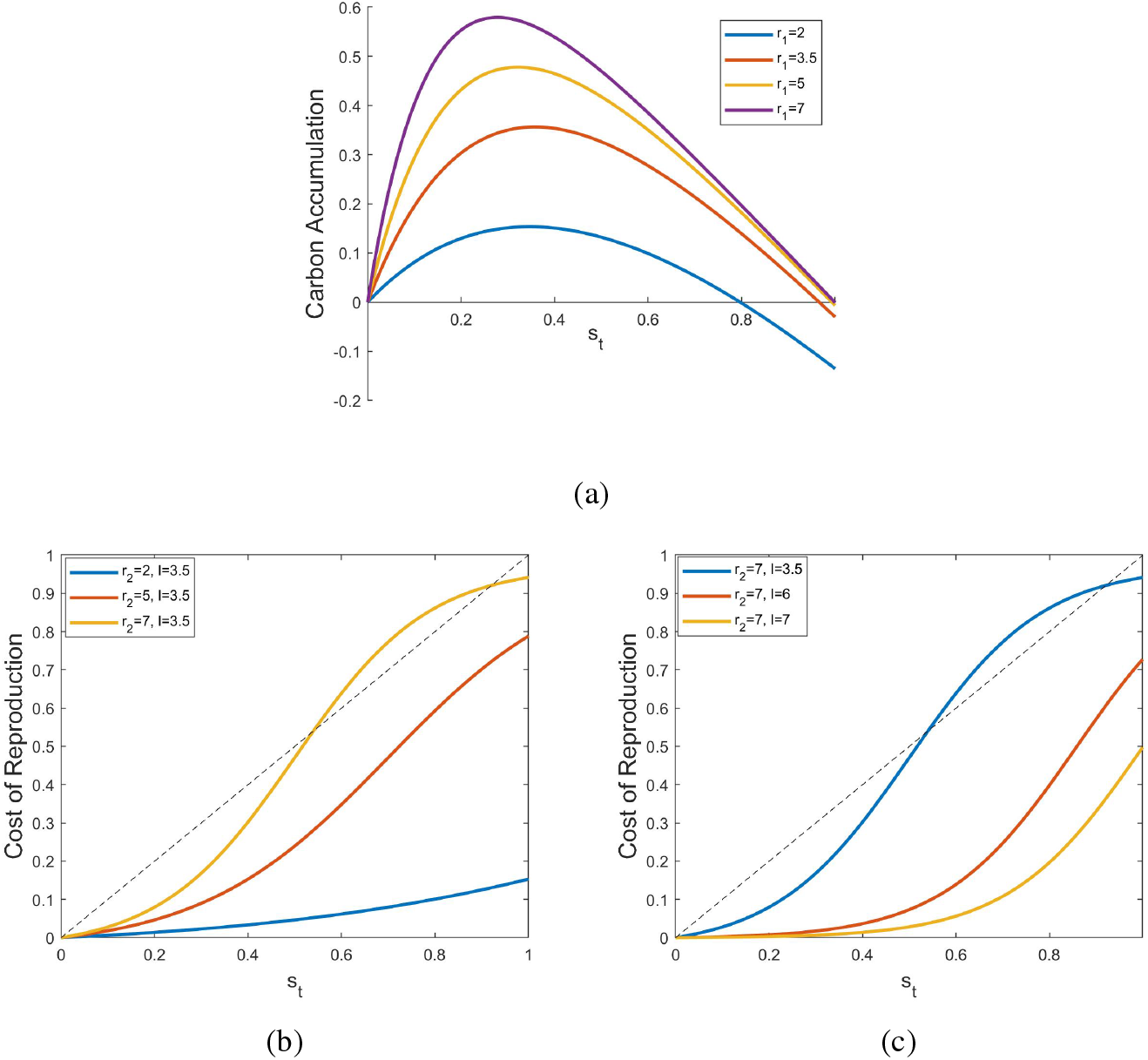
The behavior of a) nondimensionalized Carbon Accumulation term for different values of *r*_1_, b) and c) nondimensionalized Cost of Reproduction for different values of *r*_2_ when *l* = 3.5, and different values of *l* when *r*_2_ = 7, respectively.

### Shifted Sigmoid Function

The cost of reproduction is modeled by a vertically shifted sigmoid function. A sigmoid function allows for low production when the current carbon level is low. Also, the second term (vertical shift) ensures that when *S*_*t*_ = 0, there is no reproduction, and therefore no cost.

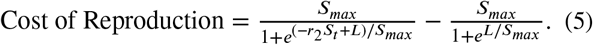

In equation 5, *r*_2_ is the tree’s reproductive investment and *L*/*r*_2_ (=*L*\*) is the threshold that controls the level of resource needed to trigger high yield. We can nondimensionalize equation 5 by defining 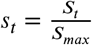 and 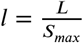. Figures 2b and 2c show the behavior of the nondimensionalized versions of the Cost of Reproduction for different values of the efficiency rate (*r*_2_) and the threshold (*l*).

As it can be seen in Figure 2a, there are values of *r*_1_ for which the Carbon Accumulation term becomes negative. Also, for some values of *r*_2_ and *l*, the curves in Figures 2b and 2c cross the diagonal line which indicates that the Cost of Reproduction exceeds the current resource levels (*s*_*t*_). For the model to be meaningful, both of these conditions must be avoided. This can be done by imposing constraints on the model and defining acceptable ranges of parameters. Section 3.2 addresses this issue in detail.

Finally, we can write the nondimensionalized model as,

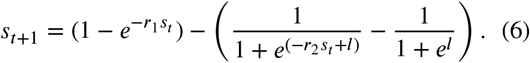

While equation 6 models the dynamics of a tree’s carbon level, the amount of nut production is the observable that is actually measured for each tree. Since nut production is the main sink of the tree’s carbon resources during reproduction, it can be taken to be proportional to the cost of reproduction. We use 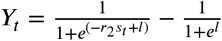 to denote the nondimension alized yield of a tree at time t to also study the dynamics of the observable of the system.

### 3.1. Bifurcation Analysis

In this section we study the behavior of the model for different values of efficiency rates and a fixed value of *l*. For simplicity, we choose the accumulation efficiency and reproductive investment rates (*r*_1_ and *r*_2_) to always be equal. Therefore, we will have *r*_1_ = *r*_2_ = *r*. This will simplify the model to a one-dimensional, single parameter map, many examples of which have been extensively explored (Strogatz, 1994; Devaney, 2003; Feigenbaum, 1980).

Figure 3a and 3b show the orbit diagram of the model and the tree’s yield (*Y*_*t*_) respectively, when *l* = 7. Qualitatively, the orbit diagrams are similar to orbit diagrams of one dimensional unimodal maps with one parameter, like the quadratic map. The model has a stable fixed point for *r* ≲ 6:8. At *r* ≲ 6:8 the first period-doubling bifurcation happens. For 6:85 ≲ *r* ≲ 8:6 the model shows a 2-cycle behavior (the range of the parameter where the alternate bearing behavior can be modeled). At *r* ≈ 8.6 a second period-doubling bifurcation happens and the system switches to a 4-cycle oscillation. Next period-doubling bifurcation happens at *r* ≈ 9.1 followed by a cascade of period-doubling bifurcations that leads to chaos. However, like the orbit diagrams of other unimodal maps, the chaotic regime is interrupted by small windows of cyclic behavior. Figure 3c shows the Lyaponov exponent as a function of the parameter *r*. For the range of the parameter values where *λ* < 0 the system has a stable fixed point or a cyclic attractor. When *λ* → −∞, the attractor is superstable. Period-doubling bifurcations happen when *λ* = 0. For *λ* > 0 the trajectories diverge exponentially which is a signature of a chaotic regime.

**Figure 3:**
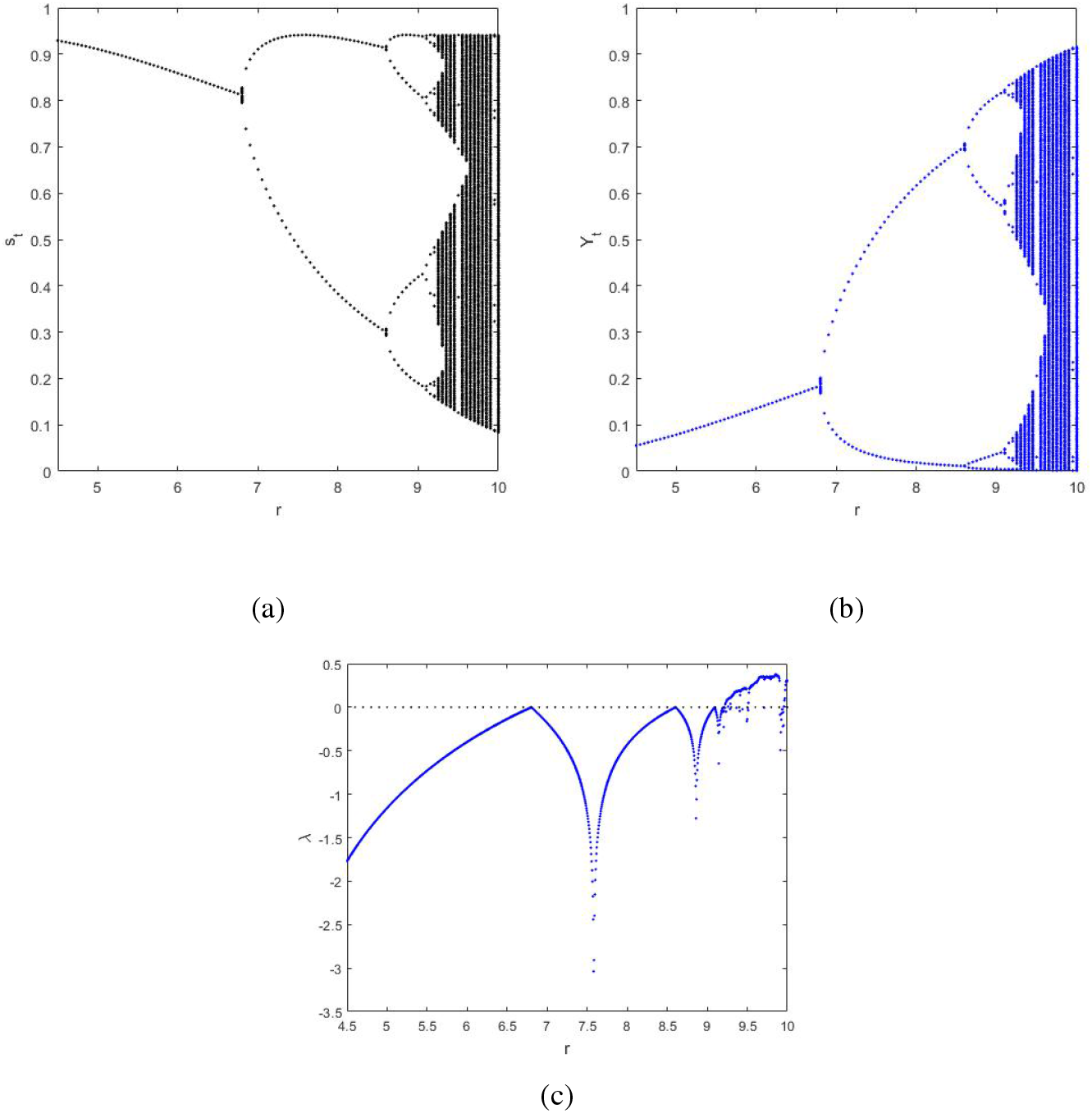
a) The orbit diagram of the model for *r*_1_ = *r*_2_ = *r* and *l* = 7, b) the orbit diagram of the tree’s yield *Y*_*t*_, and c) the corresponding Lyaponov exponent (*λ*) as a function of *r*. For values of *r* where *λ* > 0 the system is in the chaotic regime.

Figure 4 shows the trajectories for different values of *r* belonging to different regimes, again with *l* = 7. For *r* = 6, the model relaxes to a stable fixed point (Figure 4a) and the tree maintains a fixed carbon level and constant yield. For *r* = 7.5, the carbon level, and therefore the production, show a period-2 oscillation (Figure 4b). Figure 4c shows the model’s stable period-4 solution for *r* = 8.8. When *r* = 9.8 the system is in the chaotic regime (Figure 4d).

**Figure 4:**
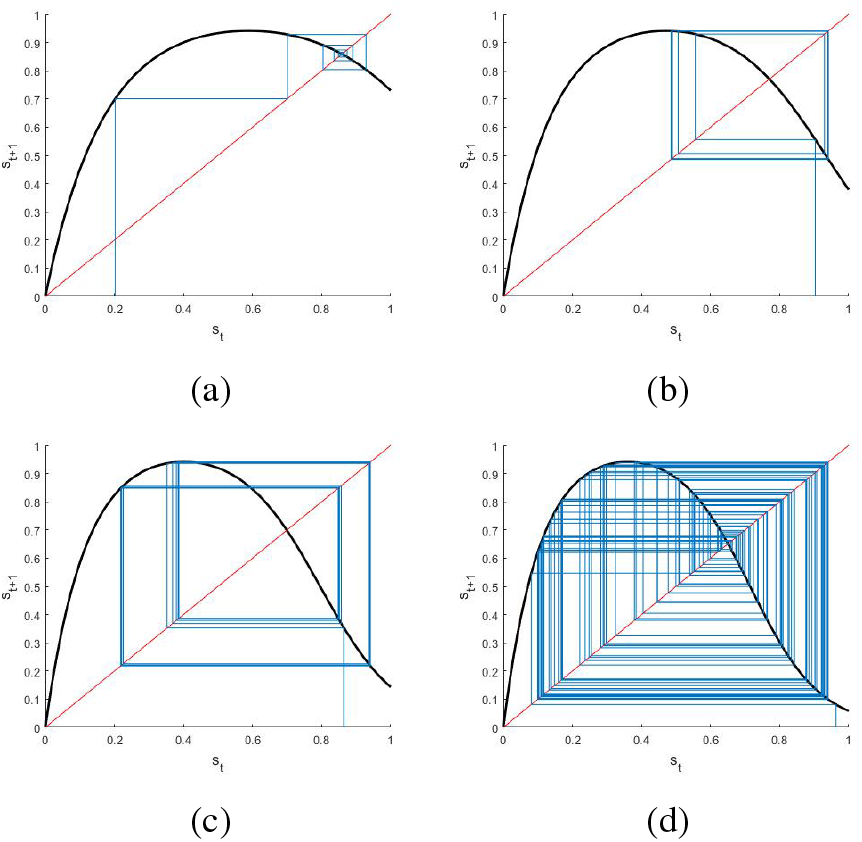
The trajectories for different values of *r*. a) *r* = 6, the model relaxes to a stable fixed point, b) *r* = 7.5, it shows a stable period-two oscillation, c) *r* = 8.8, the model has a period-four solution, d) *r* = 9.8, the systems is in the chaotic regime. In all panels *l* = 7.

The analysis in this section has also been performed for different values of *l*. As we change *l*, the locations of the period-doubling bifurcations and the width of the chaotic windows change, but the behavior of the model stays qualitatively the same.

### 3.2. Constraints on the Model

As we briefly mentioned in section 3, for a range of values of *r*_1_, the Carbon Accumulation term becomes negative. Also, for some combinations of *r*_2_ and *l* the Cost of Reproduction exceeds the current resource levels (*s*_*t*_). To avoid this and for the model to be biologically meaningful, we have to determine the acceptable range of values for the model’s parameters.

#### 3.2.1. Carbon Accumulation

The Carbon Accumulation process should always result in the increase of current carbon levels. This means that the result of equation 4 should always be greater than zero when the current carbon levels are below the maximum capacity (i.e. *S*_*t*_ < *S*_*max*_ or *s*_*t*_ < 1). As *S*_*t*_ → *S*_*max*_ (*s*_*t*_ → 1), the Carbon Accumulation function should approach zero. In other words, *S*_*t*_ = *S*_*max*_ (*s*_*t*_ = 1) should be the stable fixed point of the model when the reproduction is turned off. This means that the solution of equation 7 (Carbon Accumulation = 0) should be *s*\* = 1.

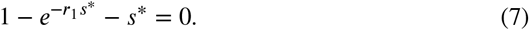

In Figure 2a, *s*\* is where each curve crosses the horizontal axis. As we can see, for any finite values of *r*_1_, the solution to the equation 7 is less than 1. This means for *s*\* < *s*_*t*_ ≤ 1 the result of the Carbon Accumulation function is nega^*t*^ tive. As it can be seen in Figure 2a, as *r*_1_ becomes larger, *s*\* gets closer to 1 and the result of the Carbon Accumulation function remains positive for a larger range of *s*_*t*_. Our goal is to find a lower bound for *r*_1_ (let’s call it *r*_*min*_) so that for values of *r*_1_ greater than *r*_*min*_ the corresponding *s*\* is sufficiently close to 1. We can write *s*\* = 1 − *δ*, where *δ* is the tolerance that controls the proximity of *s*\* to 1. To determine the *r*_*min*_ as a function of *δ*, in equation 7, we substitute *r*_1_ with *r*_*min*_ and *s*\* with 1 − *δ*. we can write,

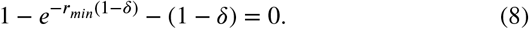

Solving for *r*_*min*_, we obtain a lower bound for *r*_1_ as a function of *δ* (i.e. *r*_1_ ≥ *r*_*min*_(*δ*)), where 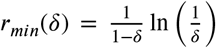. For any value of *r*_*i*_ greater than *r*_*min*_ we can accecpt that Carbon Accumulation term remains positive for *s*_*t*_ ∈ (0, 1 − *δ*) ≈ (0, 1). For the rest of this manuscript, we choose *δ* = 0.01 and study the behavior of the model for *r*_1_ ≥ 4.65.

#### 3.2.2. Cost of Reproduction

A tree’s intensity of flowering and nut production depends on the current level of its carbon storage. A tree will never draw more carbon to flower and reproduce than what is available in its reservoir. In the language of our model, the Cost of Reproduction (equation 5) cannot exceed the current carbon levels. In terms of the density of carbon levels, *s*_*t*_, it means:

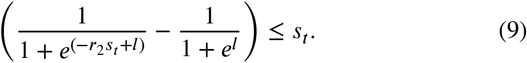

As presented in Figures 2b and 2c, for some combinations of *r*_2_ and *l*, the above condition is not met. To find the acceptable (*r*_2_*, l*) pairs, for which the condition is satisfied, we solved equation 9 numerically. The shaded area in Figure 5 shows the acceptable pairs of (*r*_2_*, l*) for which the cost of reproduction does not exceed the current carbon levels.

**Figure 5:**
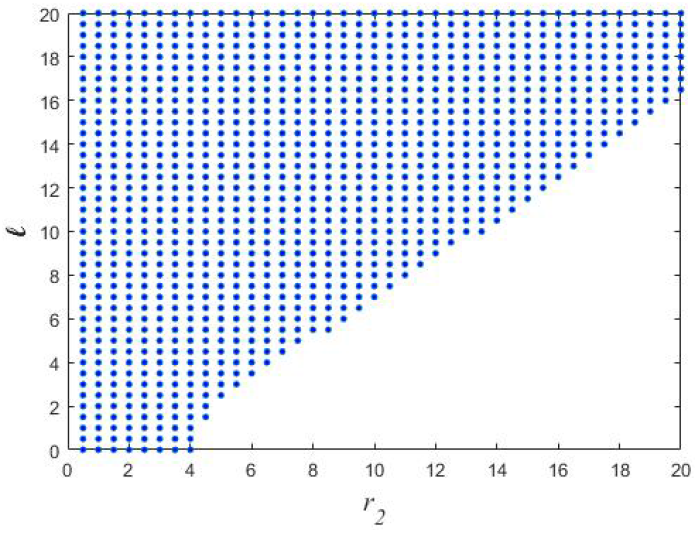
The area covered by blue dots shows the combinations of (*r*_2_, *l*) for which the condition in equation 9 is satisfied and the Cost of Reproduction stays below the current carbon levels.

## 4. The Role of Environmental Variation

The stochastic effect of environmental variation plays an portant role both in photosynthesis (the carbon accumulation process) and reproduction. Factors like the amount of CO_2_, the intensity of radiant energy, and the temperature affect the process of photosynthesis (Marshall and Biscoe, 1980). On the other hand, the amount of precipitation and the temperature during the reproduction season affect flowering or nut production. We incorporate environmental variability into the model by adding two noise terms to the carbon accumulation efficiency and reproductive investment rates (*r*_1_ and *r*_2_ respectively),

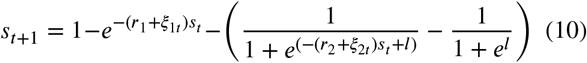

in which *ξ*_1*t*_ and *ξ*_2*t*_ are uncorrelated random variables that are independently drawn from a normal distribution with mean zero and variances 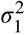 and 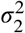, respectively.

In our simulations we choose *r*_1_ = *r*_2_ = *r* = 7 and *l* = 7 for which the system is in the two-cycle regime. We also set *σ*_1_ = *σ*_2_ = *σ*. Notice that these choices of parameters satisfy conditions set for *r*_1_, *r*_2_, and *l* as mentioned in section and shown in Figure5.While under the effect of very e noise, these conditions can be violated, for the small enough variance of the noise terms, our choices of parameters are unlikely to go beyond the acceptable range. Figure 6a shows a perfect period-two behavior of the model without noise. Figure 6b-d show the effects of noise with different strengths (variances, *σ*) on the amplitude and the phase of the oscillation.

**Figure 6:**
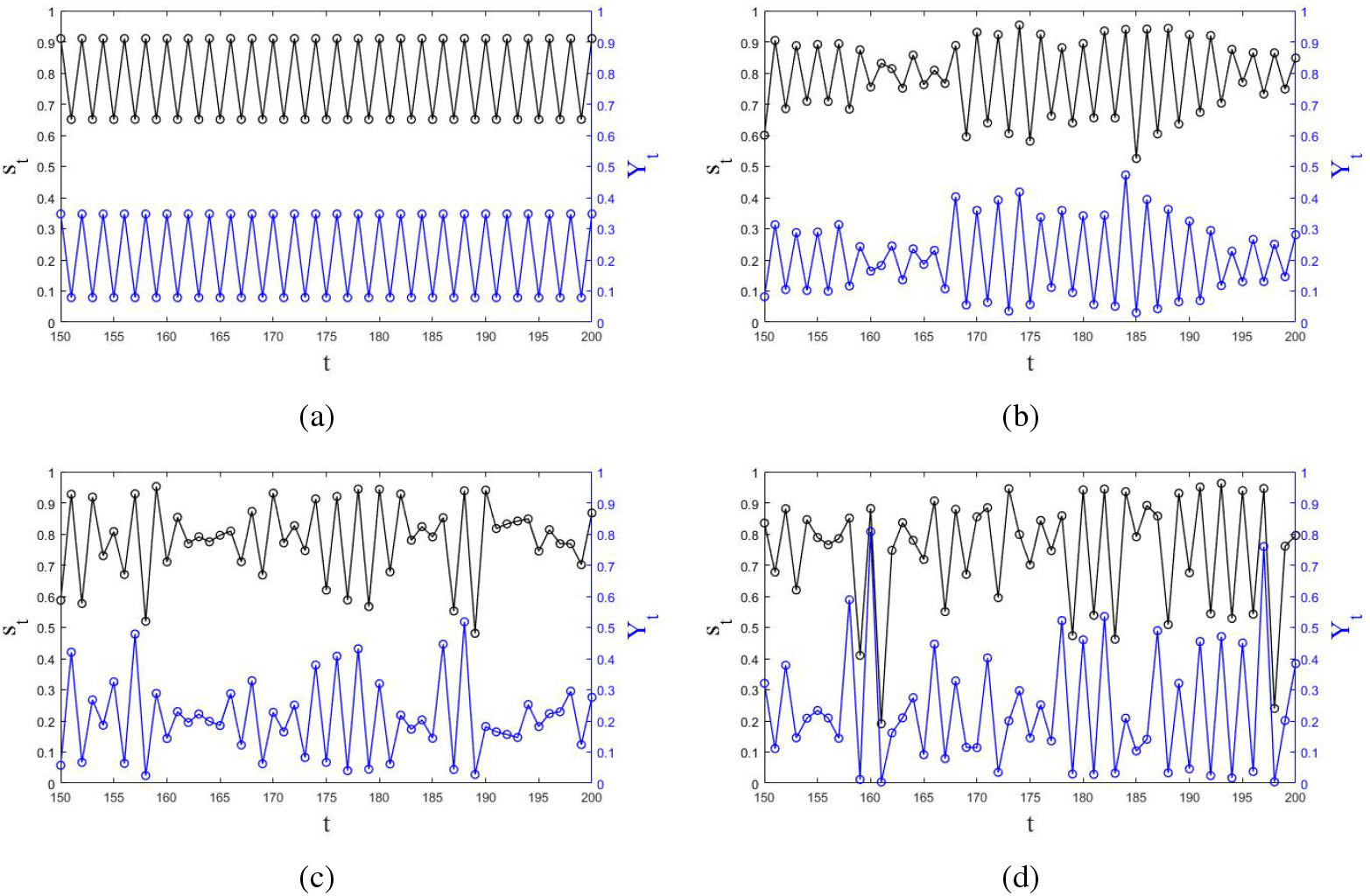
The behavior of the density of resources (black) and the yield (blue) for 50 years for *r* = 7, *l* = 7, and a) *σ* = 0 (no noise), b) *σ* = 0.25, c) *σ* = 0.5, and d) *σ* = 0.75

## 5. The Dynamics of a Two-Tree System

One of the mechanisms behind spatial synchrony, observed in the masting phenomenon, is the local interaction between trees. The trees planted in proximity to one another interact in complex ways including exchanging their carbon through root grafts (Klein et al., 2016). Grafting is known as direct interaction or diffusive coupling (Prasad et al., 2017). Trees also interact through pollination via external agents (e.g. birds, insects, and wind). This process is considered an indirect interaction and usually implemented in the form of mean-field coupling (Iwasa and Satake, 2004; Prasad et al., 2017). In dioecious plants, pollen distribution is provided by male trees while flowering and reproduction are done by female trees. Therefore, pollination cannot be considered as the mechanism behind the interaction among female trees. Instead, root grafting (direct coupling) should be considered as the method of local interaction. The numerical simulations of the Resource Budget Model with direct coupling for a system of two trees show that the only possible period-2 solution for the trees is the out-of-phase solution (Prasad et al., 2017). We confirmed these results by performing a stability analysis of the coupled Resource Budget Model as discussed in Appendix B. These results suggest that the Resource Budget Model cannot model the spatial synchrony observed among dioecious plants, like pistachios, for which the direct coupling is the main method of interaction.

In this section we use diffusive coupling to investigate the dynamics of a deterministic system of two coupled trees. The internal dynamics of each tree is defined by equation 6. We use 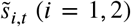 to refer to each tree’s carbon level after photosynthesis and reproduction but before exchange of resources. Each tree shares a fixed fraction of its carbon (*κ*) with its neighboring tree and receives the same fraction of the second tree’s carbon in return. The result is a net flow of carbon from one tree to another. The carbon level of each tree at the beginning of year *t* + 1 is:

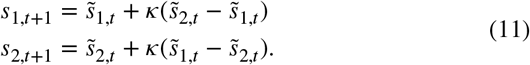

To understand the dynamics of this system, we solve uations 11 numerically to construct the orbit diagram. We assume both trees have the same internal dynamics by choosing the same carbon accumulation and reproduction efficiency rates. Also, similar to previous sections we simplify the model by setting the carbon accumulation efficiency and reproductive investment rates to be equal. Therefore, the internal dynamics of both trees only depend on one parameter, *r*. We also choose *l* = 7 for both trees. To build the orbit diagram, we follow the technique used in (Hastings, 1993). We choose 20 random initial conditions for each choice of parameters *r* and *κ* to capture all stable solutions where the system is multistable.

Figure 7 shows the dependence of the system’s dynamics on parameter *r* and different values of *κ*. Similar to the coupled logistic equations discussed in (Hastings, 1993), two general categories of solutions are identified: the perfectly in-phase solutions where *s*_1,*t*_ = *s*_2,*t*_ and all the other solutions, which we refer to as out-of-phase solutions, where *s*_1,*t*_ ≠ *s*_2,*t*_. The left column in Figure 7 shows the orbit diagram of the total carbon levels (*s*_1,*t*_ + *s*_2,*t*_) for stable in-phase solutions. The right column shows the orbit diagram of the carbon levels difference (*s*_1,*t*_ − *s*_2,*t*_) when the system has a stable out-of-phase solution. For the range of parameters for which the in-phase and out-of-phase solution coexist, both solutions are shown in red. Different patterns of oscillation are observed in both in-phase and out-of-phase categories. These patterns include in-phase or out-of-phase period-2, period-3, period-4, or higher period oscillations, as well as chaotic behavior. When the carbon exchange between the two trees is weak (e.g. *κ* = 0.05), as shown in Figure 7a and 7b, the out-of-phase solutions are more prevalent and the inphase solutions are mostly observed when the two trees are in fixed point or oscillatory regimes with different periods. As the interaction becomes stronger, the in-phase solutions appear for a wider range of parameter *r* and chaotic in-phase solutions are more commonly observed. For a strong enough *κ* (e.g. *κ* = 0.2) the trees predominantly stay in-phase while out-of-phase solutions are observed for *r* ≥ 9.7. The similarity between Figures 7e and 3a suggests that, in this case, the system behaves mostly like a single unit system.

**Figure 7:**
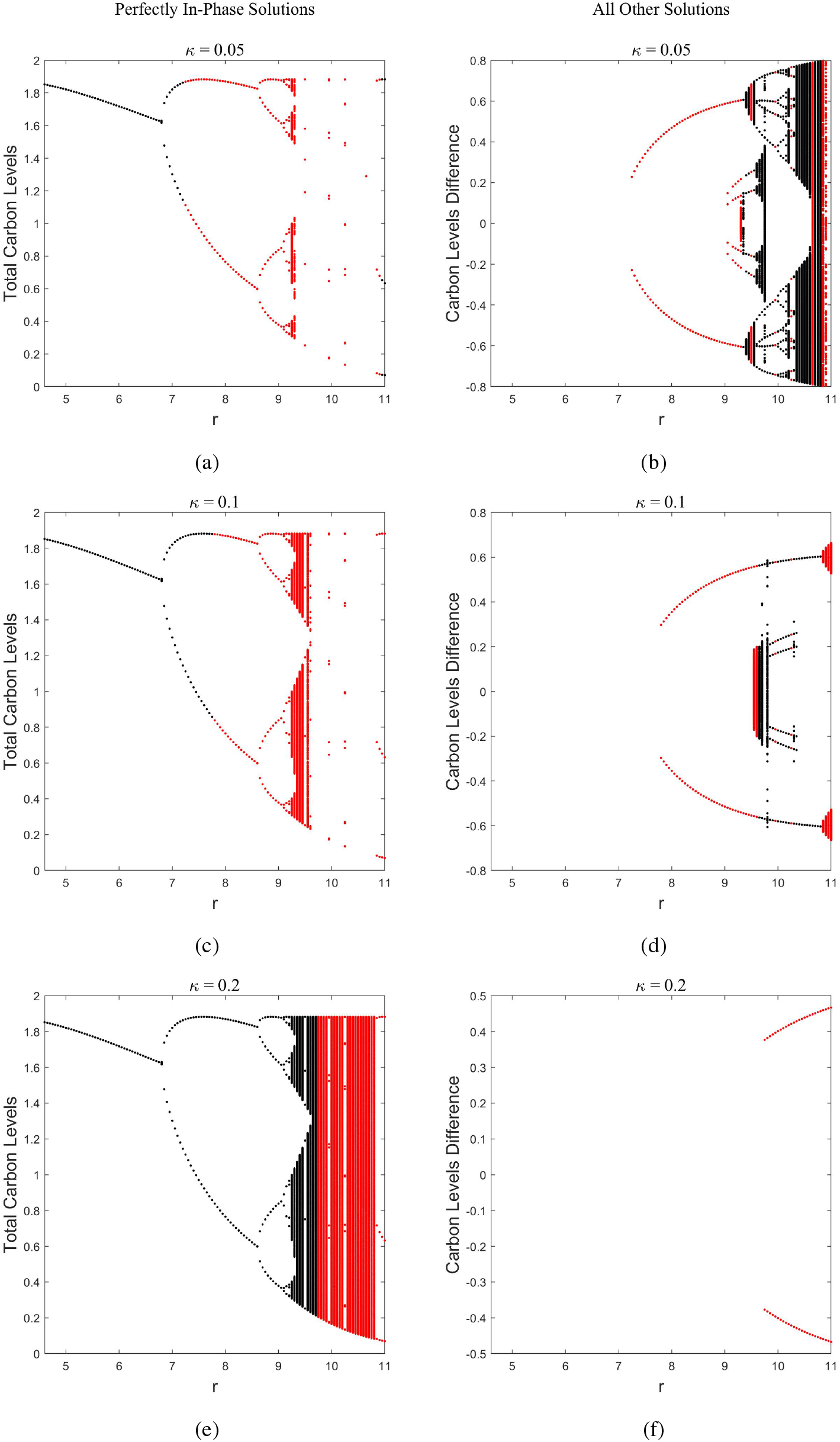
Left column: orbit diagrams of the total carbon levels when *s*_1,*t*_ = *s*_2,*t*_ (in-phase solutions), right column: orbit diagrams of the carbon levels difference when *s*_1,*t*_ ≠ *s*_2_ (all other solutions), for different values of *κ*. The red areas in all the figures indicate the coexistence of in-phase and out-of-phase solutions for that parameter value.

As we mentioned above, the category of in-phase solutions include a variety of periodic and chaotic oscillations that emerge for different values of parameter *r*. Figure 8 compares the basin of attraction of some of these solutions. To obtain Figures 8c-f, we scan the entire phase space of (*s*_1,0_, *s*_2,0_), with increment of 0.005, to find the initial conditions that relax to an in-phase attractor for *κ* = 0.1 and a given value of *r*. Figures 8c-j are color coded to match the markers lines in Figures 8a and 8b. We choose values of *r* for which the in-phase and out-of-phase solutions coexist. The in-phase solutions studied in Figure 8 are in the form of period-2 oscillation for *r* = 8 (Figure 8c and 8g), chaotic for *r* = 9.4 (Figure 8d and 8h), period-10 for *r* = 9.95 (Figure 8e and 8i), and period-3 when *r* = 10.9 (Figure 8f and 8j). The general patterns of the basins of attraction are similar for different values of *r* (different patterns of in-phase solution), however, the density and the distribution of points differ, which can provide hints toward the prevalence of the attractor in the phase space. For example, the high density and the uniform distribution of points in Figure 8c indicates that the period-2 attractor has a higher probability of emerging when *r* = 8 compared to the period-3 attractor (Figure 8f) when *r* = 10.9 which has a nonuniform basin with lower density areas.

**Figure 8:**
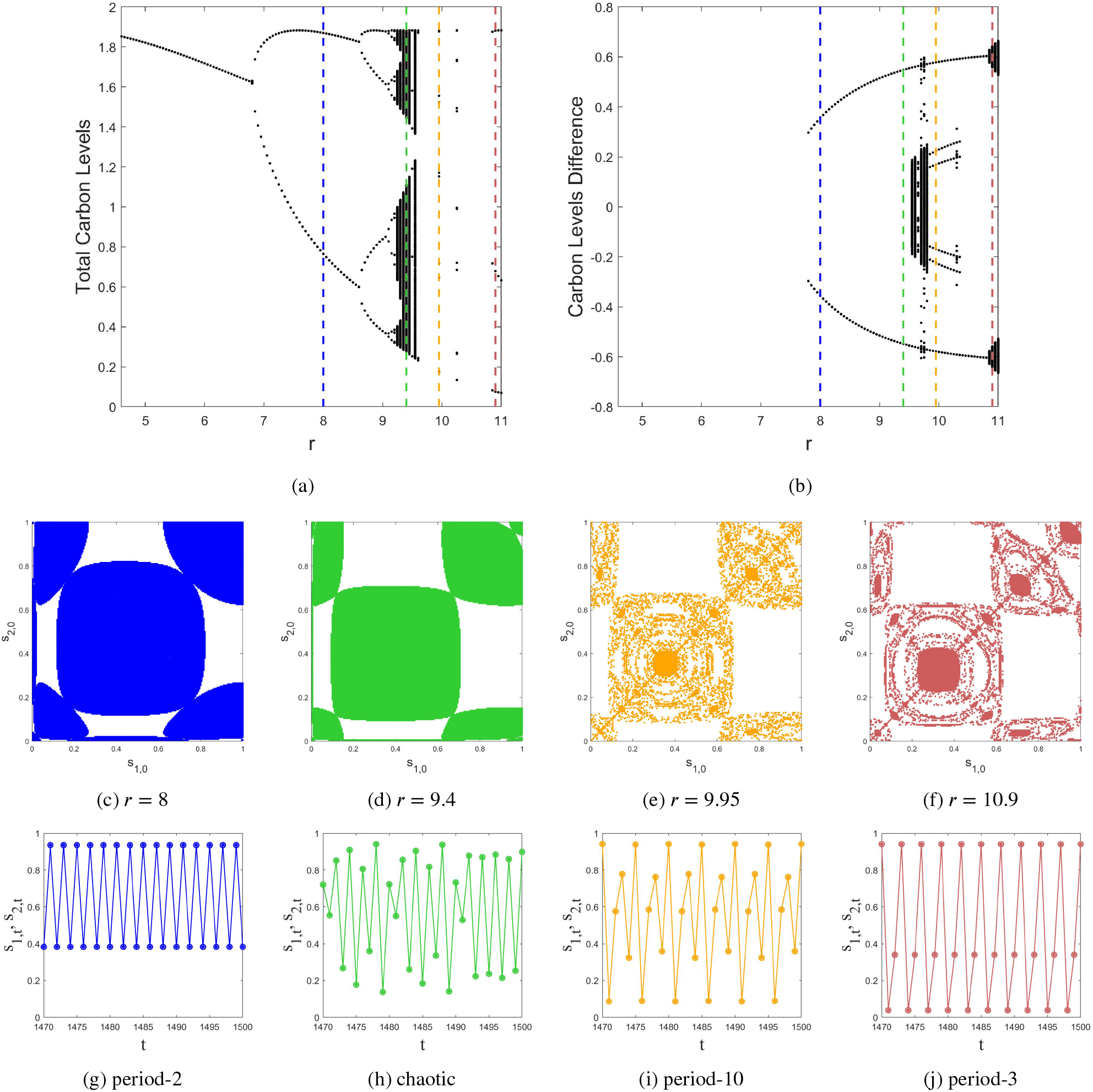
(a) and (b) The orbit diagram of in-phase (*s*_1_ = *s*_2_) and out-of-phase (*s*_1_ ≠ *s*_2_) solutions for *κ* = 0.1. (b)-(f) The basin of attraction for the in-phase solutions for different values of *r* and different patterns of oscillation (color coded to match the marker lines on (a) and (b)). (g)-(j) The corresponding pattern of in-phase oscillations for *s*_1,*t*_, *s*_2,*t*_, and *s*_1,*t*_ + *s*_2,*t*_.

## 6. Discussion

We developed a new model for the alternate bearing phenomenon. Alternate bearing is defined as the variability of the production in many types of plants in a biennial manner. Similar to the Resource Budget Model, our new model is based on the balance between generating and storing carbohydrate during photosynthesis and consuming it through flowering and nut production. We considered two biologically motivated criteria: 1) the limited capacity of each plant to store carbohydrate, and 2) low but non-vanishing yield during OFF-years (when the resource level is low). The limited capacity of the resource tank was also considered in the generalized Resource Budget Model proposed by Ye and Sakai (2016). There are multiple mathematical functions that can satisfy the above conditions. In each case different constraints should be applied to keep the model biologically meaningful. Therefore, the model can be written in different mathematical forms while the qualitative dynamics of the model remains robust.

As it was observed in the experimental data in Ye et al. (2008), the return map of the fruit production shows a hump-shaped curve. Ye and Sakai reproduced this behavior by replacing the linear relationship between resource level and the cost of flowering and fruit production in the original Resource Budget Model with a Ricker-type function which introduced more parameters to the model (Ye and Sakai, 2016). We chose proper nonlinear functions, that satisfy the biologically motivated conditions mentioned above, to model the Carbon Accumulation and Cost of Reproduction in our model. As a result the return map of the resource (carbon) level and, consequently, the yield of the plant show a logistic-like curve. Therefore, unlike the Resource Budget Model, the new model for alternate bearing has stable period-2 solutions for a wide range of the model’s parameters and is well suited to model the variable yield of plants in which the two-cycle behavior is more pronounced. The structure of our model makes it possible to nondimentionalize the carbon level of the plant and, therefore, lower the number of parameters to three. Furthermore, by setting the carbon accumulation efficiency and the reproductive investment rate to be equal, we further lower the number of parameters to two which makes the model easier to analyze and manipulate.

Trees in an orchard or natural forest do not show a perfect periodic reproduction since they are subject to environmental fluctuations. These variations affect the photosynthesis and reproduction differently. To account for environmental stochasticity, we added two uncorrelated noise terms to the accumulation efficiency and the reproductive investment rates. Our analysis shows that adding stochasticity to the model affects the amplitude and phase of the oscillation of the tree’s yield which models the noisy two cycle behavior observed in alternate bearing plants.

Masting and spatial synchrony is observed among alternate bearing plants in orchards or natural forests. As the primary step to examine the behavior of our model on a collective level, we analyzed the dynamics of a coupled two-tree system. One of the mechanisms behind masting is the local interaction between neighboring trees. This interaction can be direct (root grafting), indirect (pollen coupling), or a mixture of both. Since in diecious plants, the female trees cannot interact through pollen coupling, we used direct (or diffusive) coupling to model the local interaction. The numerical and stability analysis of the coupled Resource Budget Model with diffusive coupling showed that the only stable two cycle solution is the out-of-phase solution. Therefore, the Resource Budget Model cannot reproduce the spatial synchrony observed among female trees of the diecious plants for which root grating (direct coupling) is the main interaction mechanism. Our analysis shows that our new model for alternate bearing with direct coupling has stable in-phase period-2 solutions for a wide range of parameters and different values of coupling strength (*κ*). Having stable in-phase solutions can be interpreted as the primary requirement for the model to be used in studying spatial synchrony at a larger scale.

## Acknowledgement

We thank Patrick Brown and Todd Rosenstock for useful discussions. This work is supported by NSF grant DMS1840221.

## A. Stability Analysis of Period-2 Solutions in the Resource Budget Model

As it is shown in the orbit diagram of the Resource Budget Model in Figure 1, RBM has one fixed point 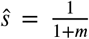 which is stable for *m* < 1. The period-2 solution, if exists, is the fixed point of *s*_*t*+2_ = *f*(*s*_*t*_), where *f*(*s*_*t*_) can be obtained by iterating the model twice. Using equation 2 we will have,

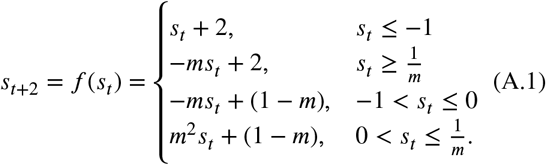

Solving 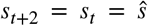, we obtain two acceptable answers from the second and fourth conditions in equation A.1,

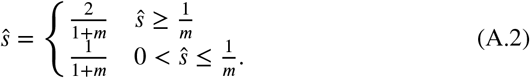

For *m* > 1 both solutions become unstable. On the other hand, if *m* < 1 the first solution becomes unacceptable since it does not satisfy its required condition that 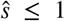. The second solution is the model’s stable fixed point and not a period-2 solution.

For *m* = 1 there is a continuum of period-2 solutions where the system oscillates between any values of 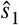 and 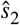, as long as 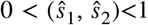, and 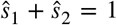. To analyze the stability of these attractors, we study the system’s response to small and large perturbations. Any small perturbation that keeps *s*_*t*_ between 0 and 1 will push the system into another period-2 attractor where the system will oscillate between two different values of 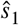 and 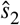. On the other hand, a large perturbation can result in *s*_*t*_ falling outside the (0, 1) range. In that case, the system will come back and settle in one of the period-2 attractors inside the continuum.

## B. Stability Analysis of the Coupled Resource Budget Model

When coupling two trees together, three scenarios can be considered:

## I) Stable fixed point

If both trees maintain the same resource levels above the eshold, we can write the model as,

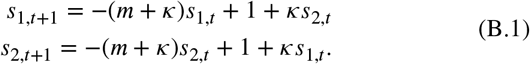

Setting *s*_1,*t*+1_ = *s*_1,*t*_ and *s*_2,*t*+1_ = *s*_2,*t*_, the fixed point of the system is 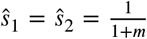. To perform the stability analysis, we find the eigenvalues of the coefficient matrix,

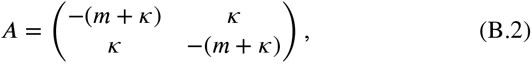

to be *λ*_1_ = −*m* and *λ*_2_ = −*m* − 2*κ*. Therefore, the fixed point of the system is only stable if (*m* + 2*κ*) < 1.

## II) Both trees oscillating between two positive values

In this case, the model is the same as equation B.1. If the trees are in-phase with the same amplitude, *s*_1,*t*_ = *s*_2,*t*_, there is no net flow of resources between the trees and the systems will be the same as two uncoupled trees. Since for an individual tree there is only a continuum of 2-period solutions when *m* = 1, the trees will stay in-phase only if they are started with equal resource levels and *m* = 1.

If *s*_1,*t*_ ≠ *s*_2,*t*_, the trees can be out of phase oscillating between two positive values. In this case, the following two conditions will be true,

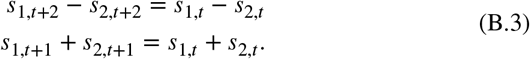

Iterating equation B.1 twice, we can write,

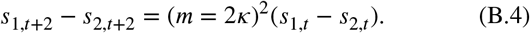

This means that the first condition in equation B.3 is satisfied if (*m* + 2*κ*) = 1. As for the second condition, we will have,

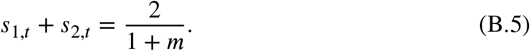

To find the period-2 solution, we assume that both trees oscillate out-of-phase between 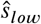 and 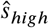 (both positive). Substituting these values in equations B.1, we will have,

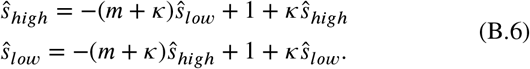

Using the criterion (*m* + 2*κ*) = 1 obtained from the first condition in equation B.3, we will have,

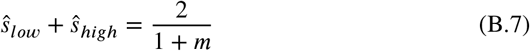

Therefore, any combination of 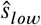 and 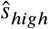 that satis equation B.7 is the solution of system.

## III) Both trees oscillating between the same positive and negative values

In this case the model can be written as,

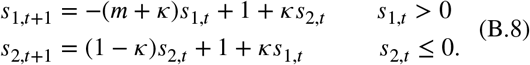

The second iteration will be,

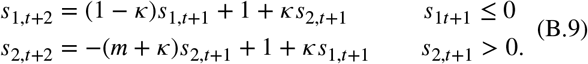

We define,

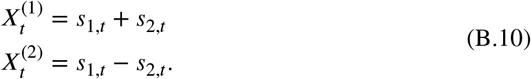

According to the conditions described in equation B.3, for the out-of-phase oscillation, we can write,

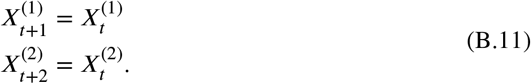

Applying the first and second iteration of the model in equations B.11,

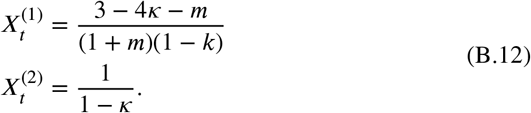

From here, we can obtain 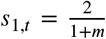 and 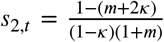. We began our discussion assuming that *s*_1,*t*_ > 0 and *s*_2,*t*_ ≤ 0. Our final result for *s*_1,*t*_ is in agreement with our initial assumption. But for *s*_2,*t*_ to be less than or equal 0, (*m* + 2*κ*) should be greater than or equal one (*m* + 2*κ*) ≥ 1).

To analyze the stability of this solution, we can analyze the stability of the fixed point of the following pair of equations,

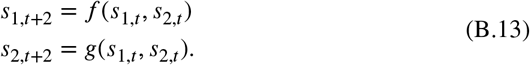

Using equations B.8 and B.9, the coefficient matrix will be,

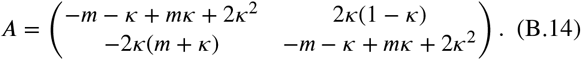

Matrix *A* has two eigenvalues, *λ*_1_ = (*κm* − *m* − *κ* + 2*κ*^2^) − 2*κ*[(*κ* + *m*)(*κ* − 1)]^(1/2)^, and *λ*_2_ = (*κm* − *m* − *κ* + 2*κ*^2^) + 2*κ*[(*κ* + *m*)(*κ* − 1)]^(1/2)^. Since *κ* < 1, *λ*_1_ and *λ*_2_ are complex conjugates. The period-2 solution is stable if |*λ*| < 1. This results in *m* < 1.

## C. An Alternate Version of the Model

As we mentioned before, there are multiple mathematical functions that satisfy the criteria discussed in section 3. As an example, we can use the Beverton-Holt function to model the Carbon Accumulation process. Therefore, the amount of carbon added to the tree during year *t* can be written as,

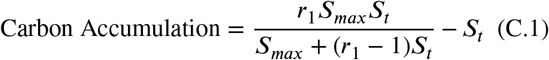

where *r*_1_ > 1 is the carbon accumulation rate, and *S*_*max*_ is the maximum capacity of the tree to accumulate carbon. We can nondimensionalize equation C.1 by defining 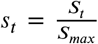 and multiplying both sides by 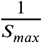. We will have,

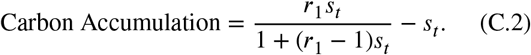

**Figure C.1:**
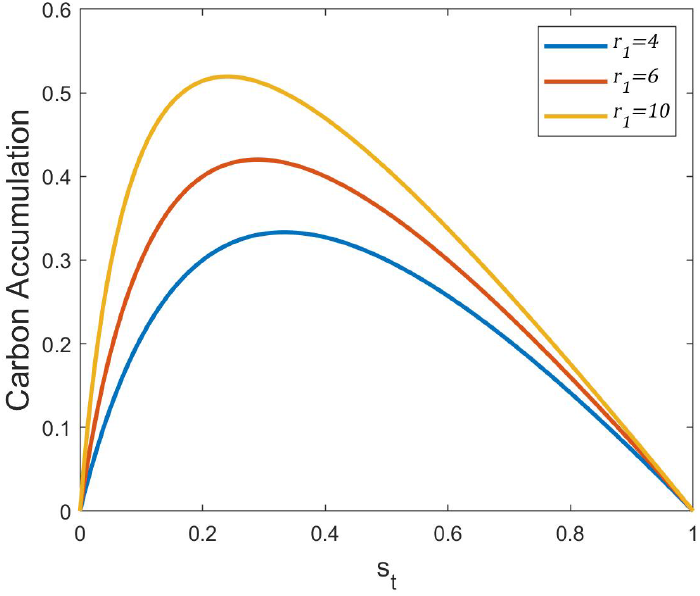
The orbit diagram of the alternate version of the model (equation C.3).

**Figure C.2:**
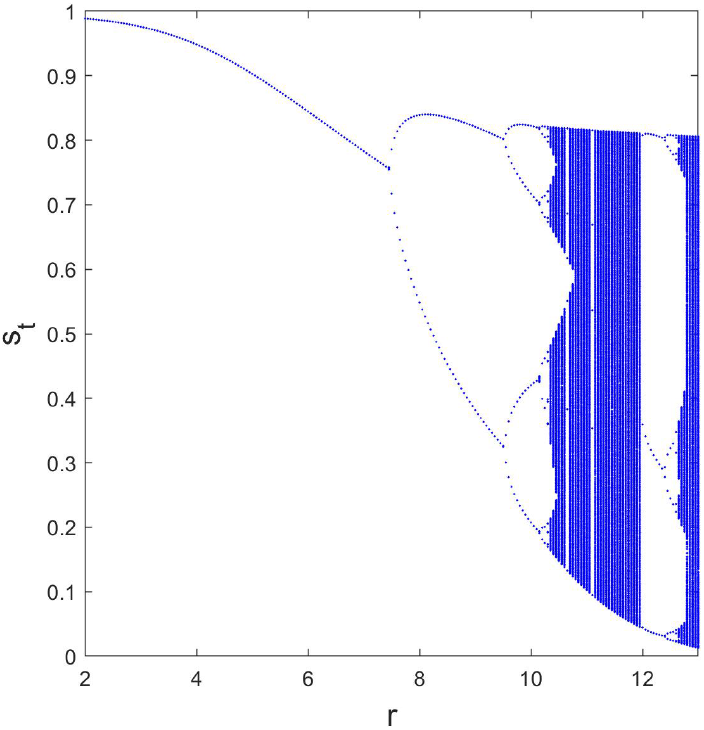
The behavior of the alternative Carbon Accumulation function (equation C.2) for different values of efficiency rate.

Figure C.1 shows the behavior of the equation C.2 for different values of *r*_1_.

We use the sigmoid function presented in equation 5 for the Cost of Reproduction. We can write the model as,

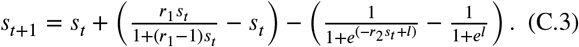

As we can see in Figure C.2, the orbit diagram of this version of the model is similar to the orbit diagram of the version proposed in the main text (Figure 3a). This is an affirmation of the robustness of the model.

